# Structural insight into tanapoxvirus mediated inhibition of apoptosis

**DOI:** 10.1101/2020.01.21.914671

**Authors:** Chathura D. Suraweera, Mohd Ishtiaq Anasir, Srishti Chugh, Airah Javorsky, Rachael E. Impey, Mohammad Hasan Zadeh, Tatiana P. Soares da Costa, Mark G. Hinds, Marc Kvansakul

**Author notes:** Centre for Virus and Vaccine Research, Sunway University, Malaysia (Mohd Ishtiaq Anasir). Co-senior author (Mark G. Hinds, Marc Kvansakul). To whom correspondence should be addressed: MK, Department of Biochemistry & Genetics, La Trobe University, Melbourne, VIC 3086, Australia.

## Abstract

Premature programmed cell death or apoptosis of cells is a strategy utilized by multicellular organisms to counter microbial threats. Tanapoxvirus (TANV) is a large double-stranded DNA virus belonging to the *poxviridae* that causes mild Monkeypox-like infections in humans and primates. TANV encodes for a putative apoptosis inhibitory protein 16L. We show that TANV16L is able to bind to a range of peptides spanning the BH3 motif of human pro-apoptotic Bcl-2 proteins, and is able to counter growth arrest of yeast induced by human Bak and Bax. We then determined the crystal structures of TANV16L bound to three identified interactors, Bax, Bim and Puma BH3. TANV16L adopts a globular Bcl-2 fold comprising 7 α-helices, and utilizes the canonical Bcl-2 binding groove to engage pro-apoptotic host cell Bcl-2 proteins. Unexpectedly, TANV16L is able to adopt both a monomeric as well as a domain-swapped dimeric topology where the α1 helix from one protomer is swapped into a neighbouring unit. Despite adopting two different oligomeric forms, the canonical ligand binding groove in TANV16L remains unchanged from monomer to domain-swapped dimer. Our results provide a structural and mechanistic basis for tanapoxvirus mediated inhibition of host cell apoptosis, and reveal the capacity of Bcl-2 proteins to adopt differential oligomeric states whilst maintaining the canonical ligand binding groove in an unchanged state.

## Introduction

Tanapoxvirus [1] is a large double-stranded DNA virus and member of the genus yatapoxvirus, which belongs to the *poxviridae* family. Tanapoxvirus (TANV) causes mild monkeypox-like infections in humans as well as primates with symptoms that include fever and skin lesions [2]. Tanapoxvirus encodes a range of immune modulatory proteins such as TNF inhibitors [3], as well as a putative B-cell lymphoma-2 (Bcl-2) homolog [4]. Bcl-2 proteins constitute a large family of proteins that primarily control programmed cell death, or apoptosis, in higher organisms [5] and are evolutionarily ancient [6]. The family comprises both prosurvival and proapoptotic members, which are characterized by the presence of one or more of four Bcl-2 homology or BH motifs and a transmembrane anchor region [7]. The mammalian prosurvival Bcl-2 members comprise Bcl-2, Bcl-w, Bcl-xL, Mcl-1, A1 and Bcl-b, and maintain host cell survival. In contrast, proapoptotic Bcl-2 family members are subdivided into two separate groups, the multimotif executors that comprise Bak, Bax and Bok, and a second group, the BH3-only proteins, which only feature a BH3 motif and includes Bad, Bid, Bik, Bim, Bmf, Hrk, Noxa, and Puma [8]. The BH3-only proteins modulate apoptosis by neutralizing the activity of prosurvival Bcl-2 through binding a surface groove [9]. After activation, Bak and Bax oligomerize to perforate the outer mitochondrial membrane leading to inner membrane herniation [10] and subsequent release of cytochrome *c*, that triggers the formation of the apoptosome and activation of downstream caspases that dismantle the cell [11].

A number of large DNA viruses encode functional, sequence and structural homologs of Bcl-2 that promote infected host cell survival and viral proliferation [12]. Viral Bcl-2 homologs have been identified in *herpesviridae*, include those from Epstein Barr virus BHRF1 [13, 14] and Kaposi sarcoma virus KsBcl-2 [15–17]. Other major virus families that contain members encoding for pro-survival Bcl-2 proteins include the *asfarviridae* with African swine fever virus encoded A179L [18–20], and grouper iridovirus encoded GIV66 [21, 22] from the *iridoviridae*. However, the largest number of Bcl-2 homologs are found in *poxviridae* [12] such as vaccinia and variola virus F1L [23–25] and myxomavirus M11L [26–28]. Whilst structural studies have shown [18, 21, 24, 29–32] these virus encoded Bcl-2-homologs that adopt a Bcl-2 fold, there is substantial diversity with regards to which proapoptotic host Bcl-2 proteins they bind. For example, vaccinia virus F1L binds Bim, Bak and Bax only [33], whereas sheeppoxvirus SPPV14 binds Bid, Bim, Bmf, Hrk, Puma, Bak and Bax [34], and African swine fever virus A179L binds all major pro-apoptotic Bcl-2 proteins [18]. The diversity observed amongst the proapoptotic ligand binding profiles for virus encoded prosurvival Bcl-2 extends to the mechanisms of action too, for example, myxomavirus M11L primarily acts by sequestering Bak and Bax [28], whereas vaccinia virus F1L neutralizes host cell death by sequestering Bim during viral infections [23].

Tanapoxvirus encoded TANV16L is a putative homolog of deerpoxvirus DPV022 [4] (Figure 1), with which it shares 32 % sequence identity. In order to understand the putative apoptosis regulatory function of TANV16L we examined its ability to bind to peptides of host proapoptotic Bcl-2 proteins, and determined crystal structures of TANV16L bound to its interactors. We now show that TANV16L is a highly flexible Bcl-2 fold protein that is able to bind to BH3 motif peptides of host proapoptotic Bcl-2 proteins with high affinity both as a monomeric and domain-swapped dimeric form. These findings provide a mechanistic basis for tanapox mediated inhibition of apoptosis, and highlight the substantial structural flexibility in the Bcl-2 fold that allows multiple oligomeric topologies to engage proapoptotic interactors using the canonical ligand binding groove.

**Figure 1:**
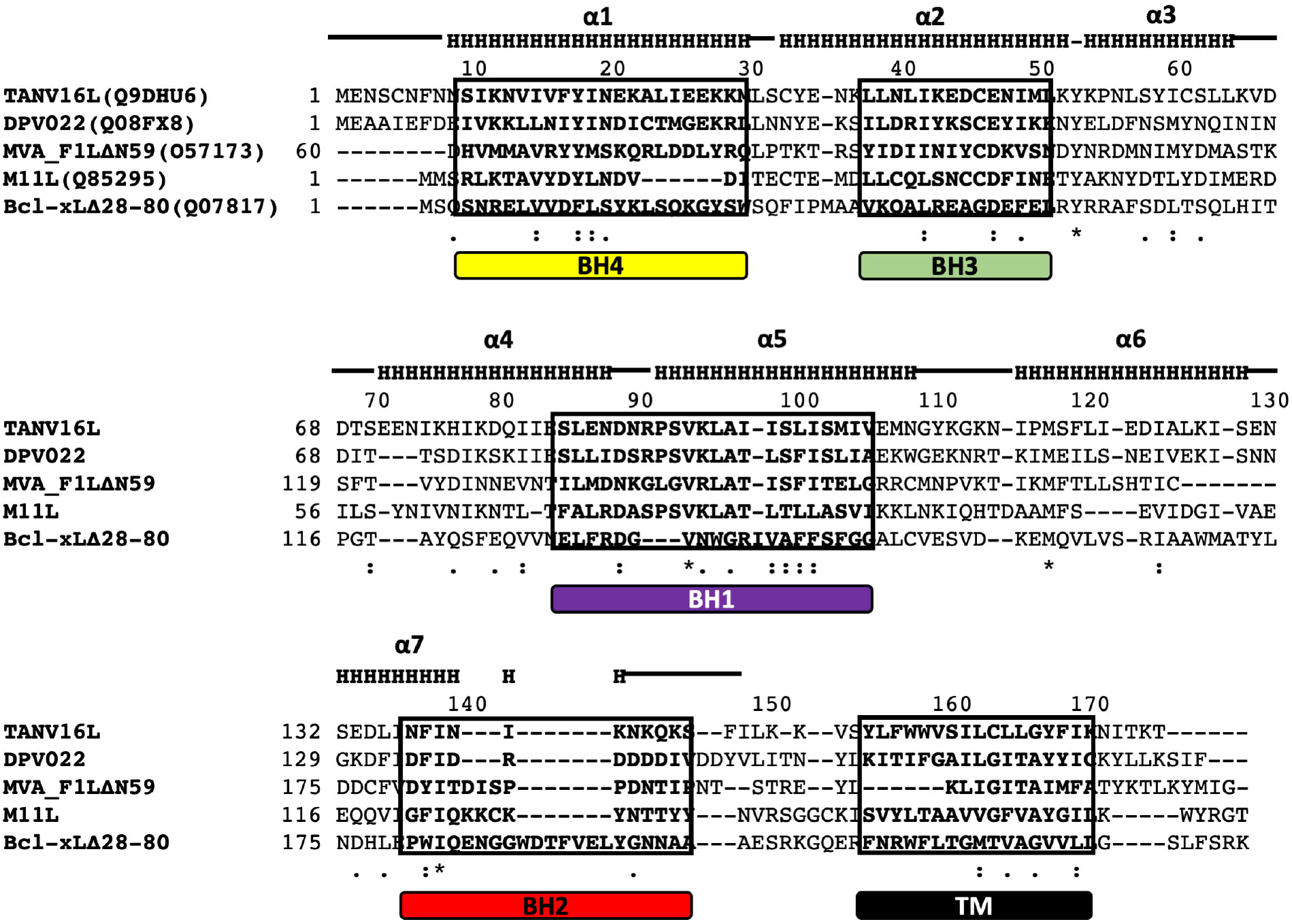
Sequence alignment of TANV16L with pro-survival Bcl-2 family members. The sequences of tanapoxvirus 16L (uniprot accession number: A7XCC0), deerpoxvirus DPV022 (uniprot accession number: Q08FX8), vaccinia virus F1L (uniprot accession number: O57173), myxomavirus M11L (uniprot accession number: Q85295) and human Bcl-x_L_ (uniprot accession number: Q07817) were aligned using MUSCLE [66]. Secondary structure elements are marked based on the crystal structure of TANV16L, and BH motifs are boxed and shown in bold [70]. The regions of helix are marked ‘H’ and unstructured loops with a bar above the sequence, conserved residues are denoted by ‘*’, with highly conservative substitutions indicated by ‘:’ and conserved substitutions indicated by ‘.’.

## Results

In order to reveal the function for TANV16L, we recombinantly expressed and purified TANV16L lacking the C-terminal 23 residues and examined its ability to bind to peptides spanning the BH3 motif of all proapoptotic human Bcl-2 proteins using isothermal titration calorimetry (ITC). TANV16L bound to a number of BH3 motif peptides with high affinity, including those from the BH3-only proteins Bim, Bid, Hrk and Puma as well as those from the multimotif executor proteins Bak and Bax (Figure 2, Table 1). We then utilized a yeast based heterologous expression system for studying functional interactions of TANV16L with Bak and Bax [13]. Consistent with our ITC data, we observed that TANV16L could directly counter Bak and Bax induced yeast growth arrest when these proteins were overexpressed in yeast (Figure 3).

**Table 1:** Interactions of TANV16L with pro-apoptotic BH3 motif peptides. All affinities were measured using isothermal titration calorimetry and KD values given in nM as a mean of three independent experiments with SD. NB no binding detectable, nd denotes that data obtained did not allow determination of affinity and thermodynamic parameters, N denotes stoichiometry of the interaction.

| Peptide | WT 16L<br>$K_D$ (nM) | $\Delta H$ | N | 16L_R90A<br>$K_D$ (nM) | $\Delta H$ | N |
| --- | --- | --- | --- | --- | --- | --- |
| Bak | $38 \pm 4$ | $-4.27 \pm 0.07$ | $0.99 \pm 0.08$ | $2325 \pm 261$ | $-4.07 \pm 0.26$ | $0.95 \pm 0.02$ |
| Bax | $70 \pm 5$ | $-4.55 \pm 0.2$ | $1.01 \pm 0.06$ | $2301 \pm 374$ | $-7.36 \pm 0.37$ | $1.03 \pm 0.03$ |
| Bok | NB | NB | NB | NB | NB | NB |
| Bad | $219 \pm 34$ | $-6.3 \pm 1.3$ | $0.95 \pm 0.07$ | $10206 \pm 1570$ | $-4.45 \pm 0.20$ | $0.87 \pm 0.06$ |
| Bid | $719 \pm 66$ | $-4.13 \pm 0.5$ | $1.02 \pm 0.07$ | nd | nd | nd |
| Bik | $1250 \pm 111$ | $-2.51 \pm 0.31$ | $0.85 \pm 0.03$ | $14356 \pm 1370$ | $-3.17 \pm 0.75$ | $1.12 \pm 0.05$ |
| Bim | $180 \pm 15$ | $-4.93 \pm 0.31$ | $0.90 \pm 0.03$ | $1766 \pm 320$ | $-6.16 \pm 0.63$ | $0.98 \pm 0.1$ |
| Bmf | $606 \pm 76$ | $-5.83 \pm 0.52$ | $0.98 \pm 0.06$ | nd | nd | nd |
| Hrk | $3220 \pm 301$ | $-5.07 \pm 0.10$ | $1.02 \pm 0.09$ | nd | nd | nd |
| Noxa | NB | NB | NB | NB | NB | NB |
| Puma | $468 \pm 47$ | $-5.87 \pm 0.15$ | $0.88 \pm 0.08$ | $911 \pm 39$ | $-1.17 \pm 0.1$ | $0.99 \pm 0.08$ |

| Peptide | 16L_K52A<br>$K_D$ (nM) | $\Delta H$ | N |
| --- | --- | --- | --- |
| Bak | $113 \pm 3$ | $-8.24 \pm 0.73$ | $1.03 \pm 0.04$ |
| Bax | $266 \pm 11$ | $-6.23 \pm 0.32$ | $1.05 \pm 0.04$ |
| Bok | NB | NB | NB |
| Bad | $5857 \pm 732$ | $-3.63 \pm 0.40$ | $0.89 \pm 0.04$ |
| Bid | $976 \pm 126$ | $-5.8 \pm 0.26$ | $1.04 \pm 0.03$ |
| Bik | $3382 \pm 85$ | $-5.67 \pm 0.11$ | $0.97 \pm 0.06$ |
| Bim | $558 \pm 27$ | $-4.61 \pm 0.29$ | $1.01 \pm 0.006$ |
| Bmf | nd | nd | nd |
| Hrk | nd | nd | nd |
| Noxa | NB | NB | NB |
| Puma | $1010 \pm 130$ | $2.95 \pm 0.15$ | $1.04 \pm 0.02$ |

**Figure 2:**
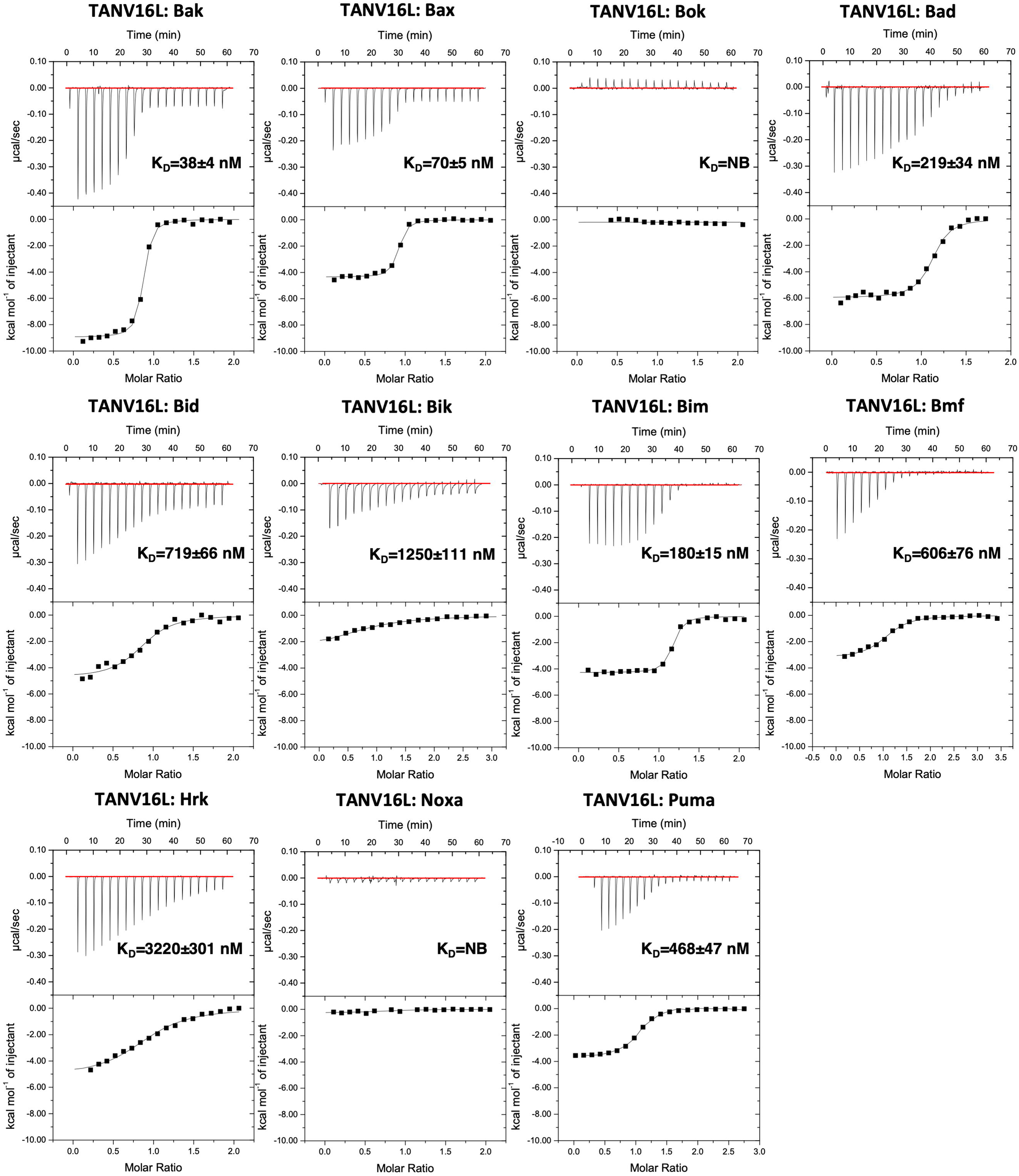
TANV16L engages a broad spectrum of BH3 motif peptides of pro-apoptotic Bcl-2 proteins. The affinities of recombinant TANV16L for BH3 motif peptides (26-mers, except for a Bid 34-mer and Bax 28-mers) were measured using ITC and the raw thermograms shown. K_D_ values (in nM) are the means of 3 experiments ± SD NB: no binding detected. The binding affinities are tabulated in Table 1.

**Figure 3:**
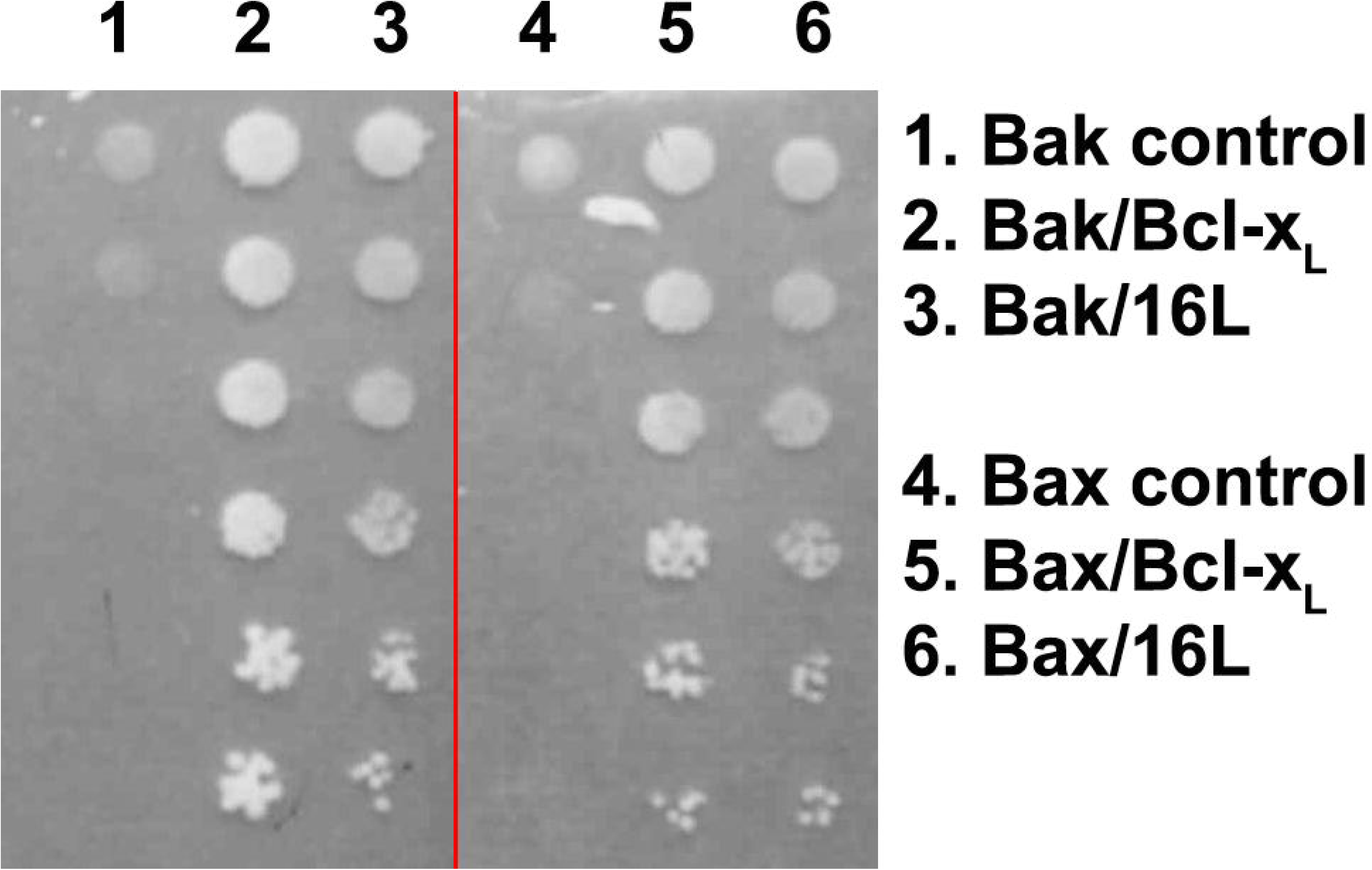
TANV16L is able to prevent Bak and Bax induced yeast growth arrest. Yeast cotransformed with constructs encoding Bax or Bak and the indicated pro-survival proteins, each under the control of an inducible (GAL) promoter, were spotted onto inducing galactose (“ON”) or repressing glucose (“OFF”) plates as 5-fold serial dilutions. Images are representative of 2 independent experiments.

To understand the structural basis for proapoptotic Bcl-2 binding by TANV16L we then determined the crystal structure of TANV16L bound to the human Bax, Bim and Puma BH3 motifs (Figure 4, Table 2). In the TANV16L:Bax BH3 complex, TANV16L adopts a globular Bcl-2 fold comprising 7 α-helices (Figure 4a). Similar to vaccinia and variola virus F1L and deerpoxvirus DPV022 TANV16L adopts a domain-swapped dimeric topology where the α1 helix from one protomer is swapped with that of a second protomer in the complex, taking up the space vacated by the matching α1 helix (Figures 4a, c). Superimposition of one chain of the TANV16L from the domain-swapped dimer TANV16L:Bax BH3 complex with the equivalent chain from the domain-swapped dimer VACV F1L from F1L:Bak BH3 [23] and the domain-swapped dimer DPV022 from DPV022:Bax BH3 [29] (Figures 4c, g) yields an rmsd of 2.3 Å (superimposed over 122 Cα atoms) and 2.1 Å (superimposed over 131 Cα atoms), respectively, whereas superimposition of the entire domain-swapped dimer of TANV16L yields rmsd values of 2.2 and 2.7 Å, respectively. The position of the TANV16L α1 within the domain-swapped dimer is identical to VACV F1L and DPV022 (Figure 4f). Similarly, TANV16L in the TANV16L:Bim BH3 complex adopts a domain swapped dimeric configuration that is identical to the TANV16L:Bax BH3 complex (Figure 4b).

**Table 2:**
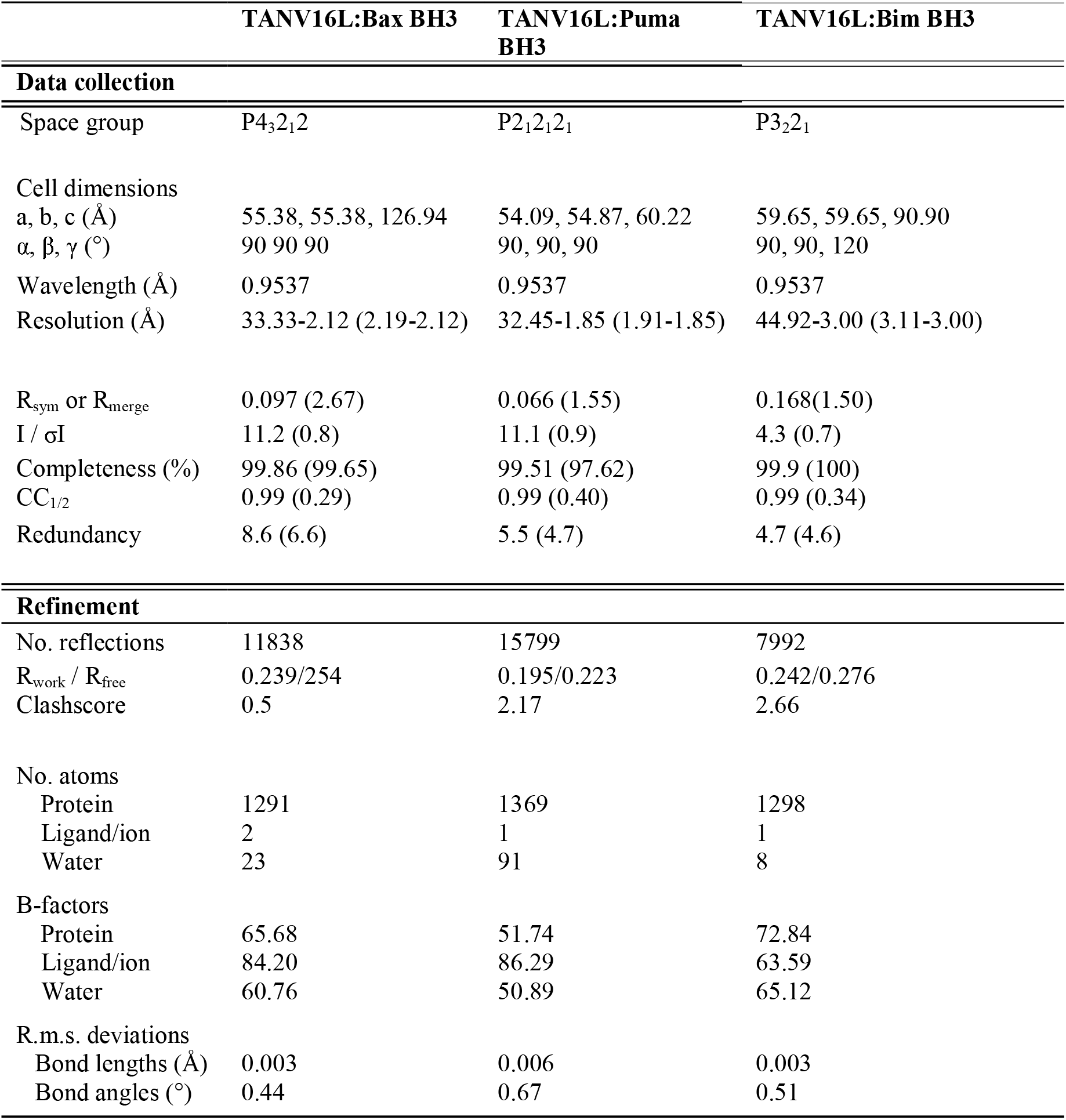
X-ray data collection and refinement statistics. Values in parentheses are for the highest resolution shell.

**Figure 4:**
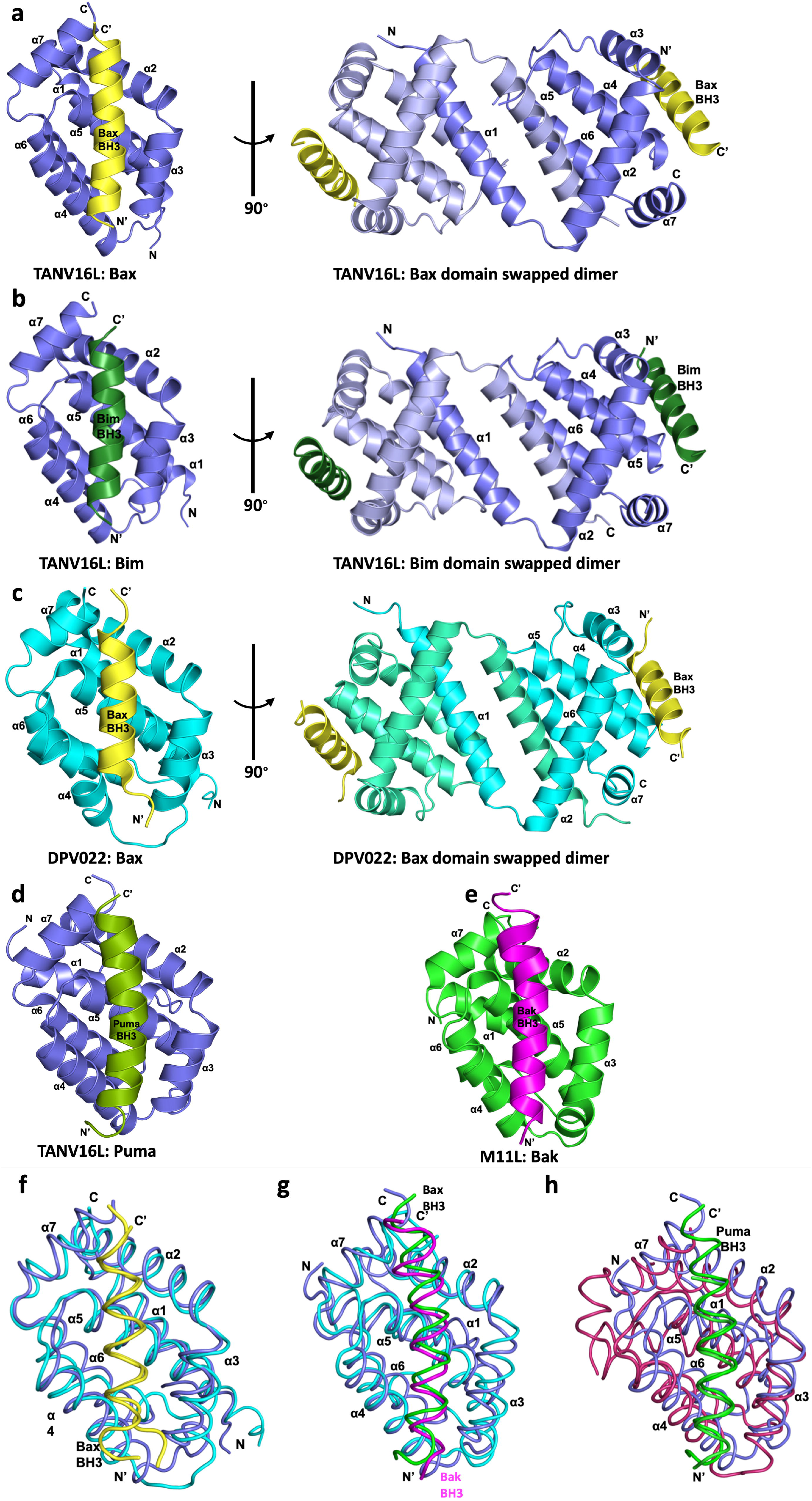
Crystal structures of TANV16L bound to Bax, Puma and Bim BH3 motifs. (**a**) TANV16L (slate) in complex with the Bax BH3 motif (yellow). TANV16L helices are labelled α1-α7. The left-hand view is of the hydrophobic binding groove of one protomer formed by helices α3-α5, the right-hand view is the domain swapped dimer viewed along the 2-fold symmetry axis between the domain-swapped α1 helices. (**b**) TANV16L (slate) in complex with the Bim BH3 domain (dark green) (**c**) DPV022 (cyan) in complex with the Bax BH3 domain (yellow) [29]. The view is as in (a). (**d**) TANV16L (slate) in complex with the Puma BH3 domain (olive) **(e)** Myxoma virus M11L (green) in complex with Bak BH3 (magenta). All structures were aligned using DALI pairwise alignment [35] and the view is as in (a). The view in the right-hand panels in (b) and (c) are as in (a). **(f)** Superimposition of the Cα backbone of TANV16L:Bax BH3 (slate and yellow) with DPV022:Bax BH3 (cyan and yellow, PDB ID 4UF2). **(g)** TANV16L:Bax BH3 (slate and yellow) with DPV022:Bak BH3 (cyan and magenta, PDB ID 4UF1) **(h)** TANV16L:Puma BH3 (slate and green) (green) with human Mcl-1:Puma BH3 (raspberry and green, PDB ID 6QFM), the most similar structure to that of TANV16L:Puma BH3 identified using Dali. All views are into the canonical ligand binding groove formed by helices α2-5. Superimpositions were generated using pairwise Dali alignment [35].

Unexpectedly, TANV16L in the TANV16L: Puma BH3 complex adopts a monomeric Bcl-2 fold, where the α1 helix is folded back into the side of the globular Bcl-2 fold (Figure 4d). A DALI analysis [35] indicated that the closest homolog in the PDB is myxomavirus M11L (Figure 4e, PDB ID 2JBX [28]) with an rmsd of 2.1 Å (over 111 Cα), whereas the closest mammalian Bcl-2 structure is human Mcl-1 (PDB ID 5FC4 [36]) with an rmsd of 2.3 Å (over 126 Cα) (Figure 4h). In the crystal structure of TPV16L:Puma BH3, one heterodimer contacts a neighbouring one via an interface formed by helices α1 and α2 from one chain and α7 as well as Puma BH3 from a neighbouring chain (Figure 5d). PISA (protein interfaces, surface and assembly) analysis of this interface yields a complexation significance score of 0, which suggests it is a crystallographic dimer and not a functionally relevant interface.

**Figure 5:**
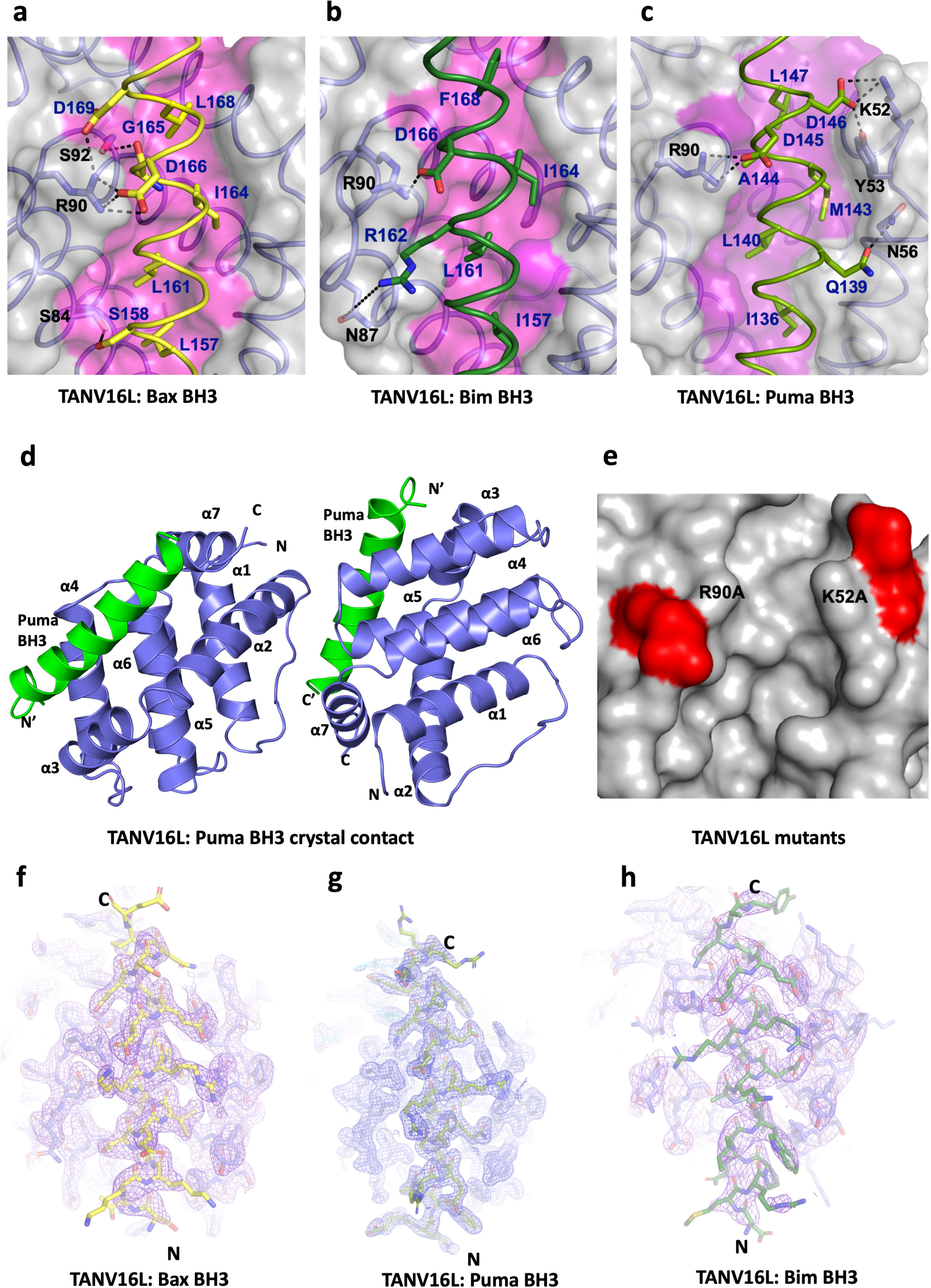
Detailed view of the TANV16L: Bax, Bim and Puma BH3 interfaces and mutation sites. (**a**) Surface depiction of the TANV16L: Bax BH3 complex. The TANV16L surface, backbone and floor of the binding groove are shown in grey and pink respectively, while Bax BH3 is shown as a yellow ribbon. The four key hydrophobic residues of Bax BH3 (L157, L161, I164 and 168L) are protruding into the binding groove, and the conserved salt-bridge formed by Bax D166 and TANV16L R90 are labeled, as well as residues involved in hydrogen bonds. (**b**) TANV16L:Bim BH3 with the surface of TANV16L is shown as in (a), and Bim BH3 is shown in dark green. The four key hydrophobic residues of Bim BH3 (I157, L161, I164 and F168) are protruding into the binding groove, and the conserved salt-bridge formed by Bim D166 and TANV16L R90 are labeled, as well as residues involved in hydrogen bonds. Interactions are denoted as black dotted lines. (**c**) TANV16L:Puma BH3 is shown as in (a), with Puma BH3 shown in olive. The four key hydrophobic residues of Puma BH3 (I136, L140, M143 and L147) are protruding into the binding groove, and the conserved salt-bridge formed by Puma D145 and TANV16L R90 are labeled, as well as residues involved in hydrogen bonds. Interactions are denoted as black dotted lines. **(d)** crystal packing of TANV16L:Puma BH3 complex. (**e)** TANV16L is shown as a grey surface, with locations of mutations used to investigate the binding site shaded in red. The views were selected for the clearest view of the groove interactions in each case. (**f**) 2Fo-Fc electron density maps of TANV16L:Bax BH3 complex interface. **(g)** 2Fo-Fc electron density maps of TANV16L:Puma BH3 complex interface. **(h)** 2Fo-Fc electron density maps of TANV16L:Bim BH3 complex interface. All maps were contoured at 1.5 σ. The N- and C-termini of the BH3 peptides are labelled and the structures aligned as in Figure 4. The view was selected for clarity of the BH3 peptide side chains.

TANV16L utilized the canonical Bcl-2 ligand binding groove formed by α2-α5 to engage BH3 motif ligands (Figure 5). In the TANV16L:Bax BH3 complex (Figure 5a), Bax residues L157, L161, I164 and L168 protrude into the four hydrophobic pockets of TANV16L. In addition, ionic interactions are observed between TANV16L R90 guanidium group and Bax D166 carboxyl as well as TANV16L R90 guanidium group and Bax D169 carboxyl group, with a further two hydrogen bonds between the TANV16L S84 hydroxyl group and the Bax S158 hydroxyl group as well as TANV16L S92 hydroxyl group and the main chain amide group of Bax G165 (Figure 5a).

In the TANV16L:Bim BH3 complex (Figure 5b), Bim residues I157, L161, I164 and F168 are used to engage the four hydrophobic pockets in TANV16L. Furthermore, there is one ionic interaction between TANV16L R90 guanidium group and Bim D166 carboxyl group and a hydrogen bond between TANV16L N87 amide and Bim R162 guanidium group. In the TANV16L: Puma BH3 complex (Figure 5c), Puma utilizes the four hydrophobic residues I136, L140, M143 and L147 to engage the four hydrophobic pockets in TANV16L. These hydrophobic interactions are supplemented by ionic interactions between TANV16L R90 guanidium group and Puma D145 carboxyl group as well as TANV16L K52 ammonium group and Puma D146 carboxyl group. Furthermore, two hydrogen bonds are found between TANV16L Y53 hydroxyl and Puma D146 carboxyl group, and between the TANV16L N56 sidechain amide and Puma Q139 sidechain carbonyl groups. Superimposition of monomeric TANV16L from the complex with Puma BH3 with one of the chains from the domain-swapped dimeric form of TANV16L from the Bax BH3 complex yields an rmsd of 1.2 Å over α2-7 (117 Cα atoms), indicating that despite the topology change from monomer to domain-swapped dimer the regions of TANV16L not involved in the domain swap remain near identical.

Since we observed TANV16L in both a monomeric and domain-swapped dimeric topology, we subjected TANV16L and some of its complexes with BH3 motif peptides to analytical ultracentrifugation (AUC). AUC was performed on TANV16L alone as well as on TANV16L bound to Bim, Bax and Puma BH3 (Figure 6). TANV16L alone revealed a mixture of monomeric and dimeric protein as well as a small amount of tetramers at concentrations ranging from 0.2-0.8 mg/mL, with a ratio of monomer:dimer of ~3.5:1. Similarly, complexes of TANV16L with Bim, Bax and Puma BH3 at 0.2 mg/mL also revealed a mixture of heterodimers and heterotetramers, with the ratio of TANV16L:Bim BH3 heterodimers vs heterotetramers being ~4:1, with TANV16L: Bax and Puma complexes displaying comparable ratios of ~3.5:1, respectively, closely matching the ratio observed for TANV16L alone (Figure 6).

**Figure 6:**
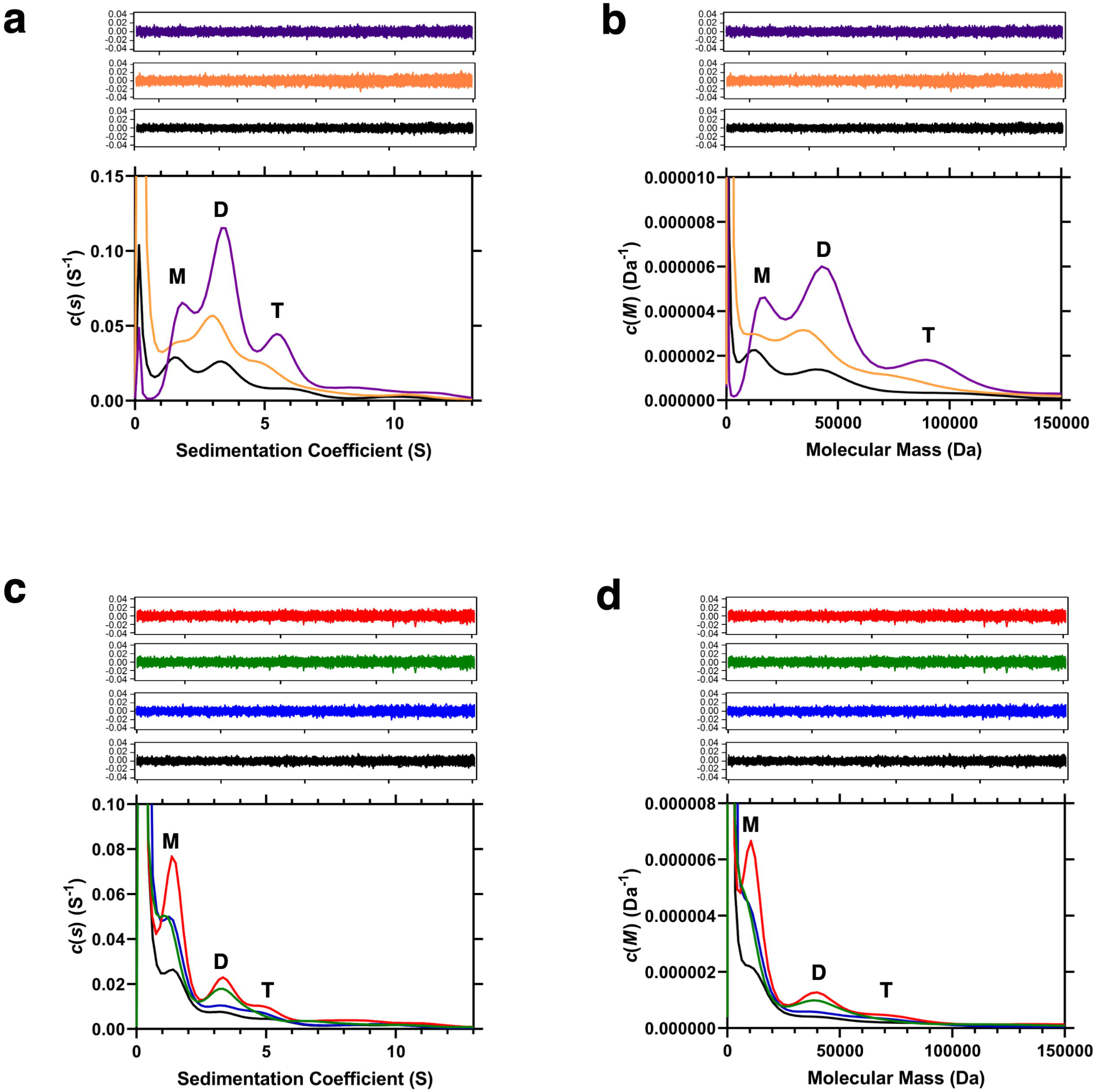
Sedimentation velocity analytical ultracentrifugation analysis of **(a,b)** TANV16L on its own at initial concentrations of 0.2 mg/ml (black), 0.4 mg/ml (orange) and 0.8 mg/ml (purple) and **(c,d)** TANV16L at an initial concentration of 0.2 mg/ml unliganded (black) and in complex with Bim (blue), Bax (green) or Puma (red). *Top panels* – Residuals resulting from the sedimentation coefficient distribution *c(s)* **(a,c)** and distribution of molar masses *c(M)* **(b,d)** best fits are plotted as a function of radial position. The residuals for the given curves are shown in the same colour above the plots. M denotes monomeric, D dimeric and T tetrameric species.

To validate the crystal structures of TANV16L bound to Bim, Bax and Puma BH3 we performed structure-guided mutagenesis (Figure 5e and analysed the mutants for their ability to bind to proapoptotic BH3 motif peptides (Table 1). Mutation of the conserved R90 in TANV16L substantially impacts on its ability to bind BH3 motif peptides, with 10-80 fold reduction in affinities for Bim, Bad, Bid, Bik as well as Bak and Bax BH3 binding, whereas binding to Hrk and Puma BH3 is only reduced ~2-fold (Figure 7). In contrast, whilst mutation of K52 to Ala also led to a 2-4 fold reduction in binding affinities for many interactors, binding to Bid BH3 was largely unaffected. Binding to Bad and Bmf BH3 was reduced by 27 and 20 fold, respectively (Figure 8).

**Figure 7:**
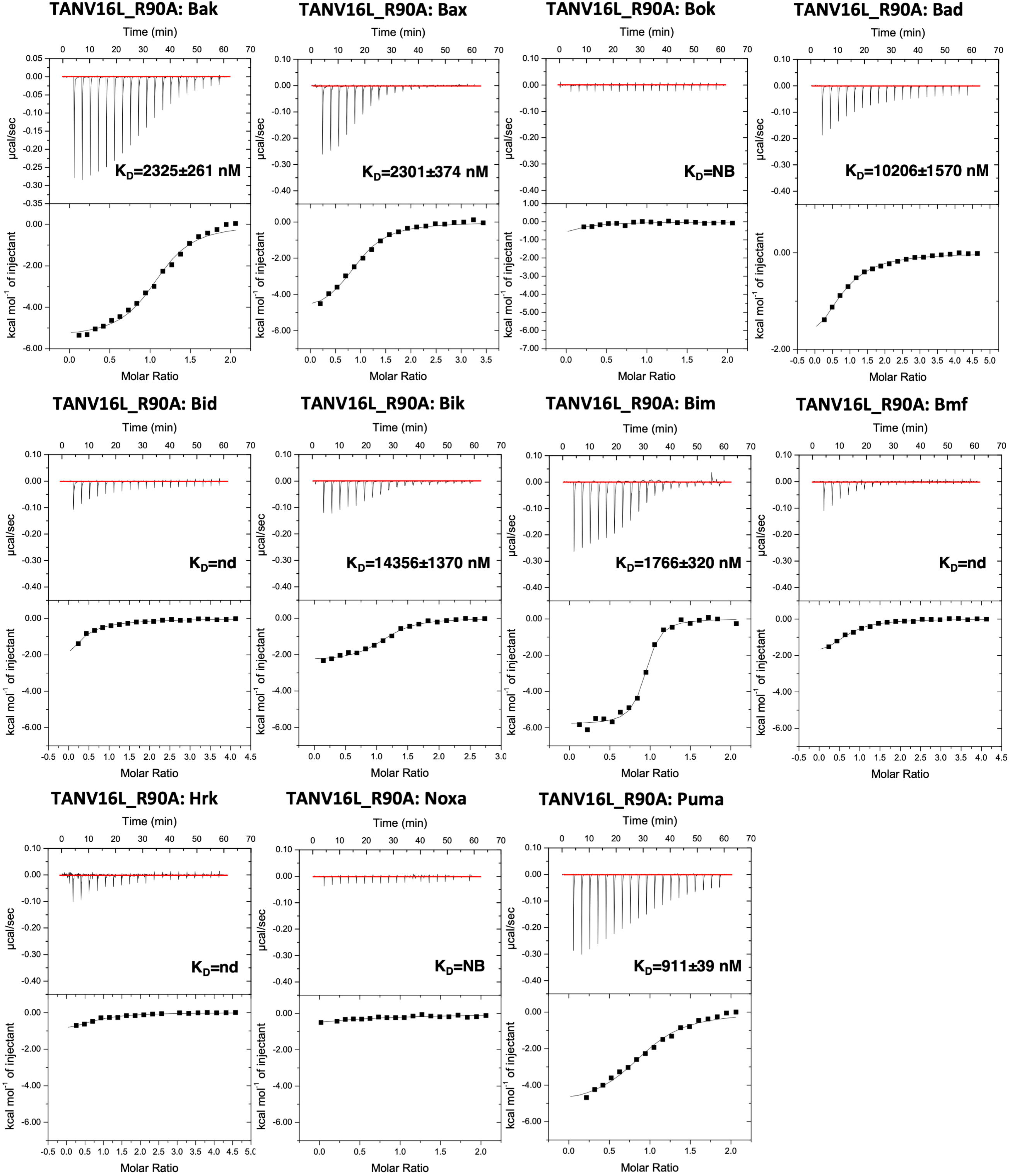
The affinities of recombinant TANV16L_R90A mutant for BH3 motif peptides (26-mers, except for a Bid 34-mer and Bax 28-mers) were measured using ITC and the raw thermograms shown. K_D_ values (in nM) are the means of 3 experiments ± SD NB: no binding detected. nd: not determined with data obtained not suitable for precise determination of affinities and thermodynamic parameters. The binding affinities are tabulated in Table 1.

**Figure 8:**
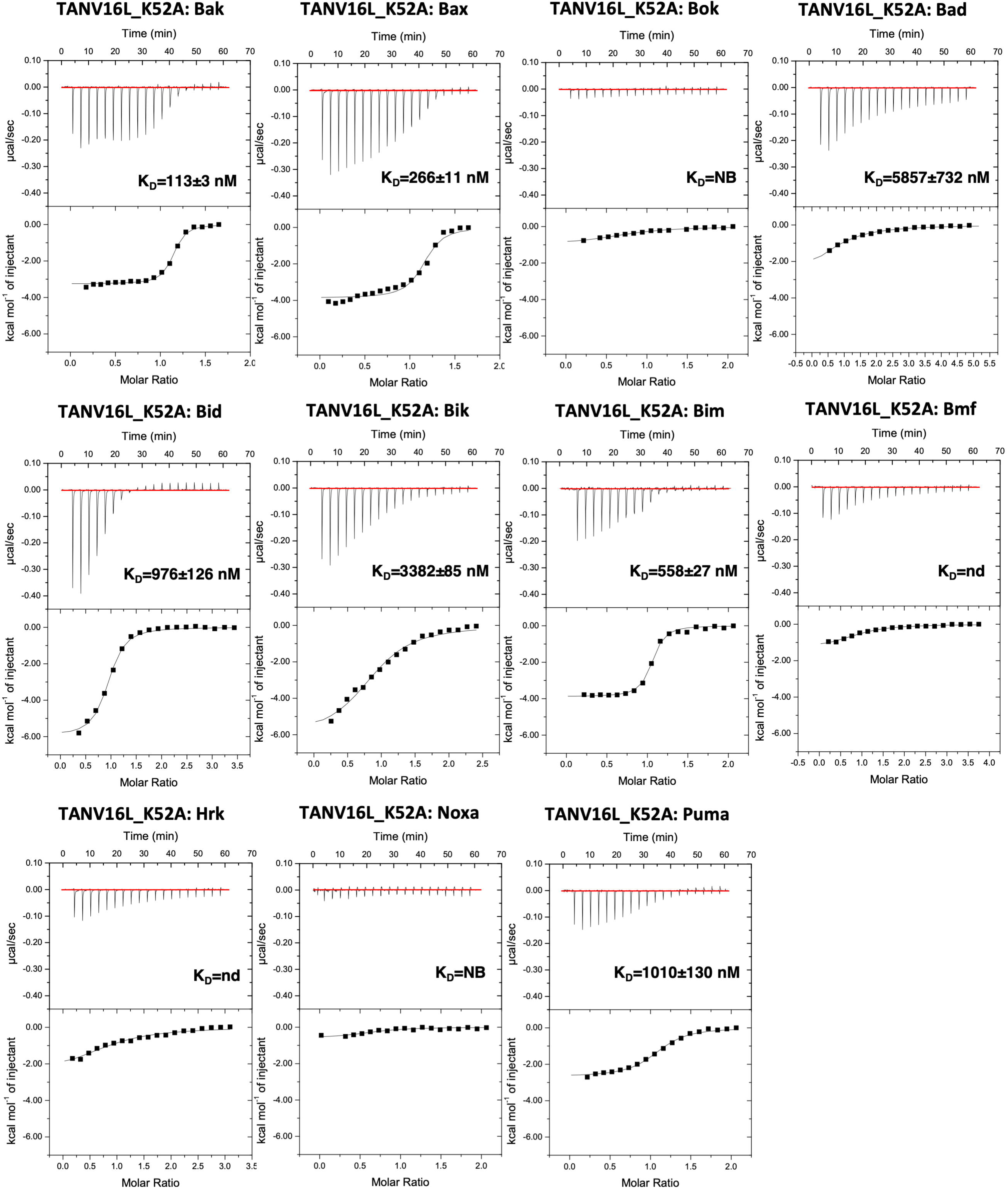
The affinities of recombinant TANV16L_K52A mutant for BH3 motif peptides (26-mers, except for a Bid 34-mer and Bax 28-mers) were measured using ITC and the raw thermograms shown. K_D_ values (in nM) are the means of 3 experiments ± SD NB: no binding detected. nd: not determined with data obtained not suitable for precise determination of affinities and thermodynamic parameters. The binding affinities are tabulated in Table 1.

## Discussion

Altruistic death of an infected cell is a potent mechanism to restrict viral infections. Viruses have evolved numerous strategies to prevent premature host cell death to establish productive infections [12]. Tanapoxvirus encodes TANV16L, a putative Bcl-2 homolog, and we now show that TANV16L adopts both a classical monomeric form as well as a domain-swapped dimer. Dimeric TANV16L is able to bind both BH3-only proteins as well as the BH3 regions of Bak and Bax. TANV16L harbours a broad proapoptotic BH3 ligand binding profile when compared to other poxvirus encoded domain-swapped pro-survival Bcl-2 proteins. The poxvirus encoded pro-survival Bcl-2 homologs VACV F1L [33], VARV F1L [24] and deerpoxvirus DPV022 [29] all have similar dimeric topologies to TANV16L dimer but feature more restricted interaction and affinity profiles. VACV F1L [33] and DPV022 [29] only bind to Bim, Bak and Bak whereas VARV F1L binds Bid, Bak and Bax [24]. In all three cases these interactions are characterized by lower affinities, in particular for Bak and Bax which bind with only micromolar affinities to VACV F1L, DPV022 F1L and DPV022. In contrast, TANV16L displays much tighter affinities including 38 nM for Bak and 70 nM for Bax BH3 (Table 1). Intriguingly, TANV16L is able to adopt both monomeric and domain-swapped dimer topologies, however the domain swap does to not impact significantly on the configuration of the canonical ligand binding groove. Superimposition of monomeric and dimeric TANV16L reveals there are virtually no differences in the respective ligand binding grooves. However, no thermodynamic analysis has been performed to examine such effects on ligand binding. When performing AUC analysis, we observed a small amount of a homotetrameric species of TANV16L in addition to monomeric and dimeric TAV16L, However, no homotetrameric structures of Bcl-2 proteins have been determined to date. Coupled with the identification of other unexpected avenues for multimerization such as a groove-in-groove dimer as shown recently for GIV66 [21] we are unable to speculate as to what a potential homotetrameric species of TAV16L may look like with regards to the overall topology and multimerization route.

A comparison of the overall dimeric structures of TANV16L with other domain swapped poxviral Bcl-2 dimers reveals that dimeric TANV16L superimposes on a VACV F1L dimer with an rmsd of 2.2 Å (over 266 Cα atoms) and on dimeric DPV022 with 2.7 Å (over 274 Cα atoms). This high level of similarity is achieved despite low overall sequence identity (8% between TANV16L and VACV F1L and 31% between TANV16L and DPV022). A detailed comparison of the interactions formed by TANV16L and DPV022 when bound to Bax reveals that the hallmark interaction between the conserved Asp from Bax BH3 with an Arg from TANV16L or DPV022 BH1 is preserved, as are hydrogen bonds between Bax Ser158 and Asp169 with TANV16L S84 or DPV022 E80. As expected, loss of the hallmark ionic interaction severely impacts TANV16L ability to bind to pro-apoptotic Bcl-2 interactors, with a TANV16L R90A mutation displaying up to 80 fold reduction in affinity (Table 1), as previously observed for mammalian prosurvival Bcl-2 [37]. Unlike in the DPV022 complex with Bax [29], the bulk of the BH3 peptides are engaged in the TANV16L ligand binding groove resulting in buried surface areas of 2343-3142 Å^2^ for the complexes with Bax, Bim and Puma, whereas in DPV022:Bax BH3 only 4 helical turns are engaged that bury 2117 Å^2^, which may contribute to the more modest affinity of the interaction. The more extensive engagement of the TANV16L binding groove with a BH3 peptide thus provides a possible rational for the substantially tighter binding observed compared with DPV022.

Virus encoded Bcl-2 proteins are not limited to domain-swapping for dimer formation (Figure 9). Vaccinia virus has been shown to encode for a suite of Bcl-2 fold proteins that modulate NF-κB signalling, with several of them adopting different dimeric topologies. N1[38], B14 and A52 [39] utilize an interface formed by helices α1 and α6, whereas A46 forms a dimer via an α4 and α6 interface [40]. In contrast, grouper iridovirus encoded GIV66 forms dimers that occlude the canonical ligand binding groove, with binding of BH3 motif peptides dissociating the dimers [21]. Domain-swapping is not limited to virus-encoded Bcl-2 proteins, and is also observed in mammalian Bcl-2 proteins. Bcl-xL has been shown to adopt domain-swapped topologies featuring either α5-α8 or α1 swaps. However, these swaps were induced by exposure to extreme pH [41] or temperature environments [42] or by truncation of the loop connecting α1 and α2 [43], respectively. In contrast, recombinantly expressed Bcl-w lacking its C-terminal tail adopts both a monomeric state as well as a domain swapped dimeric state where α3-α4 are swapped [44, 45]. Such a swap impacts the canonical ligand binding groove, and leads to differential binding affinities between monomeric and dimeric Bcl-w [45]. Domain swapping has also been shown for human proapoptotic Bak [46] and Bax [47], where the formation of an extended single helix comprising helices α5 and α6 leads to a core/latch configuration, where a core domain formed by α1-4 partners the latch domain formed by α6-α8 from a second protomer. In contrast, catfish Bax features a domain swapped α9 helix [48]. This significant number of topological variances across dimeric Bcl-2 proteins underscores the inherent flexibility in this fold, and potentially enables additional layers of functionality to modulate function in addition to direct interactions.

**Figure 9:**
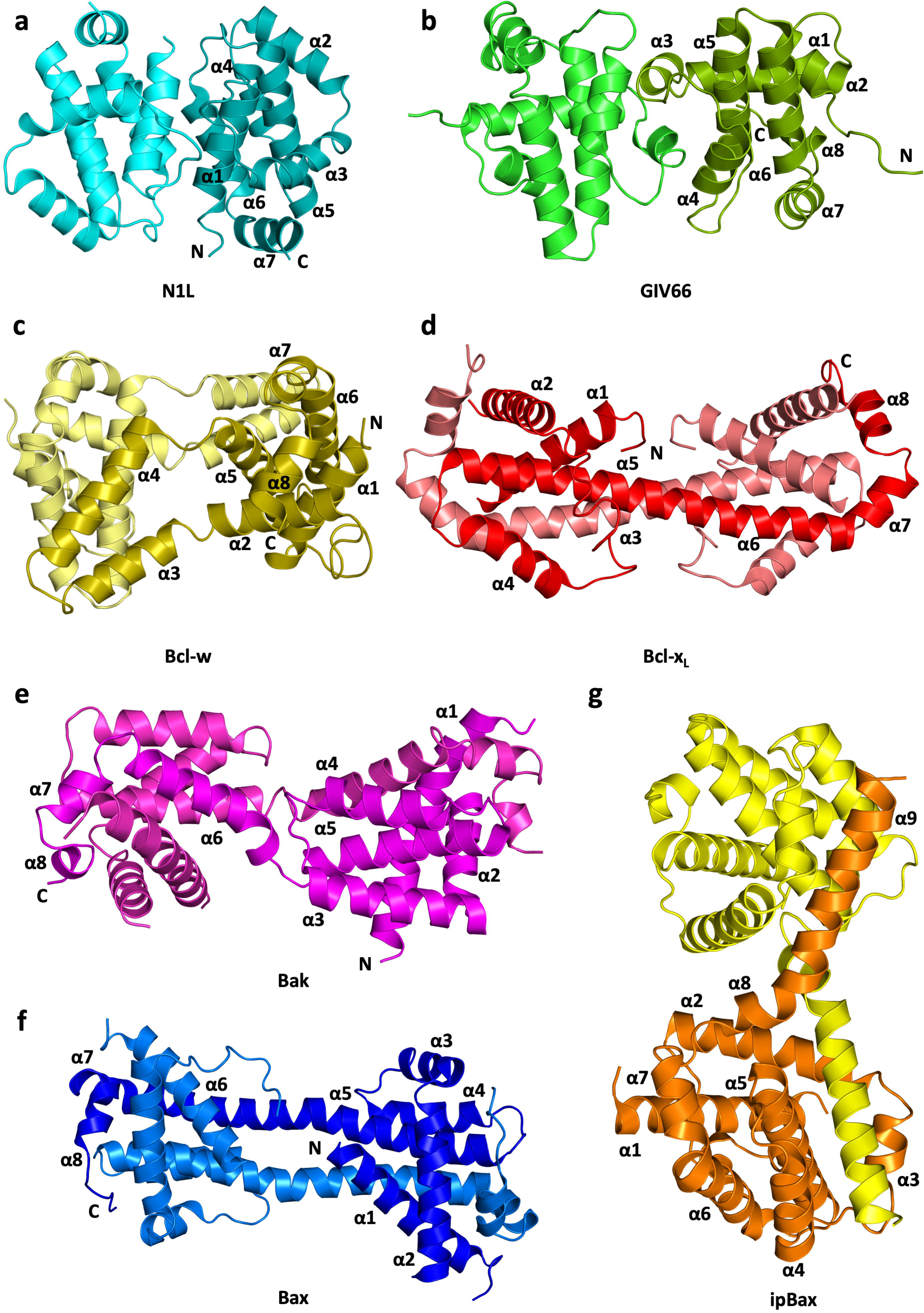
Cartoon diagrams of dimeric topologies in the Bcl-2 family **(a)** Vaccinia virus N1L homodimer (cyan, PDB ID 2UXE) [38]. **(b)** Grouper iridovirus GIV66 (green, PDB ID 5VMN) [21]. **(c)** Human Bcl-w (sand, PDB ID 2Y6W) [45]. **(d)** Human Bcl-x_L_ (red, PDB ID 1R2D) [41]. **(e)** Human Bak core-latch dimer (magenta, PDB ID 4U2U) [46]. **(f)** Human Bax core-latch dimer (blue, PDB ID 4ZIE) [47]. **(g)** Catfish Bax groove-tail dimer (yellow and orange, PDB ID 5W63) [48].

In summary, we report the biochemical and structural analysis of tanapoxvirus 16L, which revealed a broad high affinity binding profile for mammalian pro-apoptotic Bcl-2 proteins. Furthermore, our crystal structures of TANV16L bound to Bax and Puma BH3 indicate that TANV16L displays substantial structural plasticity, being able to adopt both a classical monomeric Bcl-2 fold as well as a domain-swapped dimeric Bcl-2 fold. Overall, our findings provide a mechanistic platform for dissecting the role of 16L for tanapoxvirus replication and infectivity.

## Materials and methods

### Protein expression and purification

Synthetic cDNA encoding for codon optimized wildtype TANV16L (Uniprot Accession number Q9DHU6) as well as two mutants TANV16L (K52A and R90A) lacking 23 C-terminal residues were cloned into the bacterial expression vector pGex-6p-1 (Genscript). Recombinant TANV16L was expressed in C41(DE3) cells in 2YT medium supplemented with 1 mg/ml ampicillin at 37°C in a shaking incubator until an OD_600_ of 0.6 was reached. The protein expression was induced by adding isopropyl β-D-1-thiogalactopyranoside (IPTG) to final concentration of 0.75 mM for 18 hours at 20°C. Bacterial cells were harvested by centrifugation at 5000 rpm (JLA 9.1000 rotor, Beckman Coulter Avanti J-E) for 20 min and re-suspended in 100 ml lysis buffer A (50 mM Tris pH 8.0, 300 mM NaCl and 10 mM DTT (dithiothreitol). The cells were homogenized using an Avestin EmulsiFlex homogenizer and lysed using sonication (programme 7, Fisher Scientific™ Model 705 Sonic Dismembrator) and the resultant lysate was transferred into SS34 tubes for further centrifugation at 18,000 rpm (JA-25.50 rotor, Beckman Coulter Avanti J-E) for 30 min. The supernatant was loaded onto 5 mL of glutathione sepharose 4B (GE Healthcare) equilibrated with buffer A. After sample application, the column was washed with 150 ml of buffer A and protein on-column cleavage was achieved by adding HRV 3C protease overnight at 4°C. The cleaved protein was eluted using buffer A, with the remaining protein being concentrated using a centrifugal concentrator with 3 kDa molecular weight cut-off (Amicon^®^ Ultra 15) to a final volume of 2 ml. Concentrated TANV16L was subjected to size-exclusion chromatography using a Superdex S200 increase 10/300 column mounted on an ÄKTA Pure system (GE Healthcare) equilibrated in 25 mM HEPES pH 7.5, 150 mM NaCl and 5 mM TCEP (Tris(2-carboxyethyl)phosphine hydrochloride), and fractions analysed using SDS-PAGE. The final sample purity was estimated to be greater than 95% based on SDS–PAGE analysis. Appropriate fractions were pooled and concentrated using a centrifugal concentrator with 3 kDa molecular weight cut-off (Amicon^®^ Ultra 15) to final concentration of 5.4 mg/ml.

### Analytical ultracentrifugation

Sedimentation velocity experiments were performed in a Beckman Coulter XL-A analytical ultracentrifuge as described previously [49–52]. Briefly, double sector quartz cells were loaded with 400 μl of buffer (25 mM HEPES pH 7.5, 150 mM NaCl, 5mM TCEP) and 380 μl of sample (solubilized in buffer). For the runs with apo protein, initial concentrations of 0.2 mg/ml, 0.4 mg/ml and 0.8 mg/ml were employed. For the runs with the complexes, the protein and peptide concentrations were kept at 0.2 mg/ml. The cells were loaded into an An50-Ti rotor and the experiments conducted at 25°C. Initial scans were carried out at 3,000 rpm to determine the optimal wavelength and radial positions. Final scans were performed at 40,000 rpm and data were collected continuously at 230 nm using a step size of 0.003 cm without averaging. Solvent density, solvent viscosity and estimates of the partial specific volume of *apo*-TANV16L, TANV16L:Bim, TANV16L:Puma and TANV16L:Bax. at 25°C were calculated using SEDNTERP [53]. Data were fitted using the SEDFIT software (www.analyticalultracentrifugation.com) to a continuous size-distribution model [54–56].

### Measurement of dissociation constants

Binding affinities were measured using a MicroCal iTC200 system (GE Healthcare) at 25°C using wild type TANV16L as well as two mutants TANV16L K52A and R90A in 25 mM HEPES pH 7.5, 150 mM NaCl, 5 mM TCEP. Measurements were performed at a range of different concentrations for BH3 motif peptide and TANV16L proteins, which are summarized below in Table 3. All affinity measurements were performed in triplicate. Protein concentrations were measured using a Nanodrop UV spectrophotometer (Thermo Scientific) at a wavelength of 280 nm. Peptide concentrations were calculated based on the dry peptide weight after synthesis. The BH3-motif peptides used were commercially synthesized and were purified to a final purity of 95% (GenScript) and based on the human sequences as previously described [57].

**Table 3:** Summary of protein and ligand concentrations used in ITC measurements

| **Interaction** | **Protein concentration**<br>**(μM)** | **BH3 motif peptide**<br>**concentration (μM)** |
| --- | --- | --- |
| TANV16L:Bak BH3 | 60 | 600 |
| TANV16L:Bax BH3 | 60 | 600 |
| TANV16L:Bok BH3 | 60 | 600 |
| TANV16L:Bad BH3 | 60 | 550 |
| TANV16L:Bid BH3 | 60 | 600 |
| TANV16L:Bik BH3 | 60 | 750 |
| TANV16L:Bim BH3 | 60 | 600 |
| TANV16L:Bmf BH3 | 60 | 800 |
| TANV16L:Hrk BH3 | 60 | 600 |
| TANV16L:Noxa BH3 | 60 | 600 |
| TANV16L:Puma BH3 | 60 | 700 |
| TANV16L_R90A:Bak BH3 | 60 | 600 |
| TANV16L_R90A:Bax BH3 | 60 | 800 |
| TANV16L_R90A:Bok BH3 | 60 | 600 |
| TANV16L_R90A:Bad BH3 | 60 | 950 |
| TANV16L_R90A:Bid BH3 | 60 | 900 |
| TANV16L_R90A:Bik BH3 | 60 | 700 |
| TANV16L_R90A:Bim BH3 | 60 | 600 |
| TANV16L_R90A:Bmf BH3 | 60 | 900 |
| TANV16L_R90A:Hrk BH3 | 65 | 900 |
| TANV16L_R90A:Noxa BH3 | 60 | 600 |
| TANV16L_R90A:Puma BH3 | 60 | 600 |
| TANV16L_K52A:Bak BH3 | 60 | 550 |
| TANV16L_K52A:Bax BH3 | 60 | 550 |
| TANV16L_K52A:Bok BH3 | 60 | 600 |
| TANV16L_K52A:Bad BH3 | 60 | 1000 |
| TANV16L_K52A:Bid BH3 | 60 | 800 |
| TANV16L_K52A:Bik BH3 | 60 | 650 |
| TANV16L_K52A:Bim BH3 | 60 | 600 |
| TANV16L_K52A:Bmf BH3 | 60 | 850 |
| TANV16L_K52A:Hrk BH3 | 65 | 600 |
| TANV16L_K52A:Noxa BH3 | 60 | 600 |
| TANV16L_K52A:Puma BH3 | 60 | 600 |

### Crystallization and structure determination

Crystals for TANV16L: Bax BH3, TANV16L: Puma BH3 or TANV16L: Bim BH3 complexes were obtained by mixing TANV16L with human Bax BH3 28-mer or Puma BH3 26-mer peptide into 1:1.25 molar ratio as described previously [58] and concentrated using a centrifugal concentrator with 3 kDa molecular weight cut-off (Amicon^®^ Ultra 0.5) to 5 mg/ml and concentrated protein was immediately used for crystallization trials. Initial high throughput sparse matrix screening was performed using 96 well sitting drop trays (swissic, Neuheim, Switzerland) using 200 nL of protein mixed with 200 nL of reservoir solution.

TANV16L: Bax BH3 crystals were grown by the sitting drop vapour diffusion method at 20°C in 1.0 M LiCl, 0.1 M Citrate pH 4.0, 20 % W/V PEG 6000. The crystals were flash cooled at −173°C in mother liquor supplemented with 20% ethylene glycol. Diffraction data were collected at the Australian Synchrotron MX2 beamline using an Eiger detector with an oscillation range 0.1° per frame with a wavelength of 0.9537 Å, integrated using XDS [59] and scaled using AIMLESS [60]. Molecular replacement was carried out using PHASER [61] with the previously solved structure of DPV022 (PDB ID: 4UF1 [29]) as a search model. TANV16L: Bax BH3 crystals contained one molecule of TANV16L and one Bax BH3 peptide in the asymmetric unit, with 46.3% solvent content and final TFZ and LLG values of 8.0 and 52.79 respectively. The final model of TANV16L: Bax BH3 was built manually over several cycles using Coot [62] and refined using PHENIX [63].

TANV16L: Puma BH3 crystals were grown as the TANV16L: Bax BH3 crystals and were obtained in 0.1 M Potassium thiocyanate, 30% PEG 2000MME. The crystals were flash cooled at −173°C in mother liquor. Diffraction data collection, integration and scaling were performed as described above. The molecular replacement was carried out using PHASER with the previously solved structure of TANV16L: Bax BH3 as a search model. TANV16L: Puma BH3 crystals contain one molecule of TANV16L and one Puma BH3 peptide, with 46.3% solvent content and final TFZ and LLG values of 13.2 and 133.15 respectively. The final model of TANV16L: Puma BH3 was built manually over several cycles using Coot and refined using PHENIX.

TANV16L: Bim BH3 crystals were grown similar to other two complexes as above and in 0.1 M MIB buffer pH 8.0, 25% PEG 1500. The crystals were flash cooled at −173°C in mother liquor. Diffraction data collection, integration and scaling were performed as described above. Molecular replacement was carried out using PHASER with the previously solved structure of TANV16L: Bax BH3 as a search model. TANV16L: Bim BH3 crystals contain one molecule of TANV16L and one Bim BH3 peptide, with 44.01% solvent content and final TFZ and LLG values of 15.8 and 208.64 respectively. The final model of TANV16L: Bim BH3 was built manually and refined as described above. Coordinate files have been deposited in the Protein Data Bank under the accession codes 6TPQ, 6TQQ and 6TRR. All images were generated using the PyMOL Molecular Graphics System, Version 1.8 Schrödinger, LLC. All software was accessed using the SBGrid suite [64]. All raw diffraction images were deposited on the SBGrid Data Bank [65] using their PDB accession code 6TPQ, 6TQQ and 6TRR.

### Sequence alignment and interface analysis

Sequence alignments were performed using MUSCLE [66] (https://www.ebi.ac.uk/Tools/msa/muscle/) with the default settings, and sequence identities were calculated based on the total number of conserved residues in TANV16L against the full sequence. Protein interfaces were analysed using PISA [67].

### Yeast colony assays

*Saccharomyces cerevisiae* W303α cells were co-transformed with pGALL(TRP) vector only, pGALL(TRP)-Bcl-xL, or pGALL(TRP)-TANV16L and pGALL(Leu)-Bak or pGALL(Leu)-Bax. pGALL(TRP) and pGALL(Leu) places genes under the control of a galactose inducible promoter [68]. Cells were subsequently spotted as a 5-fold serial dilution series onto medium supplemented with 2% w/v galactose (inducing, “ON”) to induces protein expression, or 2% w/v glucose (repressing, “OFF”), which prevents protein expression, as previously described [69]. Plates were incubated for 48 h at 30°C and then photographed.

## Abbreviations

(TANV): tanapoxvirus
(ITC): isothermal titration calorimetry
(Bcl-2): B-cell lymphoma 2
(BH): Bcl-2 homology

## Acknowledgements

We thank staff at the MX beamlines at the Australian Synchrotron for help with X-ray data collection. We thank the ACRF for their support of the Eiger MX detector at the Australian Synchrotron MX2 beamline and the Comprehensive Proteomics Platform at La Trobe University for core instrument support. This research was funded by the Australian Research Council (Fellowship FT130101349 to MK and DE190100806 to TPSC), National Health and Medical Research Council of Australia (CDA fellowship 637372 and Project Grant APP1007918 to MK) and La Trobe University (Scholarship to CDS, MIA, AJ and REI).

## Author Contributions

Chathura D. Suraweera: Acquisition, analysis and interpretation of data; Drafting and revising the article.

Mohd Ishtiaq Anasir: Acquisition, analysis and interpretation of data.

Airah Javorsky: Acquisition, analysis and interpretation of data;

Srishti Chugh: Acquisition, analysis and interpretation of data.

Rachael E. Impey: Acquisition, analysis and interpretation of data.

Muhammad Hasan Zadeh: Acquisition, analysis and interpretation of data.

Tatiana P. Soares da Costa: Acquisition, analysis and interpretation of data; Drafting and revising the article.

Mark G. Hinds: Conception and design; Analysis and interpretation of data; Drafting and revising the article;

Marc Kvansakul: Conception and design; Acquisition of data; Analysis and interpretation of data; Drafting and revising the article.

